# Establishment and Characterization of a *CCND1*-Rearranged Non-Mantle Cell Lymphoma Cell Line and Patient-Derived Xenograft Model

**DOI:** 10.1101/2025.08.23.671479

**Authors:** Claus-Moritz Gräf, Moritz Reese, Angela Vicente-Luque, Nicolas Mönig, Charlotte Bruzeau, Ferran Nadeu, Maria Latacz, Johanna Bihler, Jörn Meinel, Sílvia Beà, Elias Campo, Melanie Thelen, Paul J. Bröckelmann, Ron D. Jachimowicz

**Author notes:** These authors contributed equally to this work (Paul J. Bröckelmann, Ron D. Jachimowicz).

## Abstract

The pathobiology of aggressive B-cell lymphomas with *CCND1* rearrangements, distinct from Mantle Cell Lymphoma (MCL), presents a significant clinical challenge. These lymphomas are often difficult to diagnose and demonstrate resistance to standard immunochemotherapy, underscoring the urgent need for a deeper understanding of their underlying biology to develop more effective treatments. A major impediment to progress has been the lack of robust preclinical models that accurately reflect the complex genomics and clinical behavior of this disease. Here we directly address this critical gap by reporting the establishment and in-depth characterization of the first patient-derived cell line and a corresponding systemic patient-derived xenograft (PDX) model of a *CCND1*-rearranged, non-MCL lymphoma with a rapidly fatal clinical course. Through a comprehensive multi-omics approach, we demonstrate that these novel *in vitro* and *in vivo* models faithfully recapitulate the primary tumor’s unique immunophenotype, its intricate genetic and transcriptional landscape, and its intrinsic resistance to conventional therapeutic agents.

## To the Editor

Accurate diagnosis of distinct lymphoma subtypes critically informs contemporary therapeutic strategies, which increasingly incorporate targeted agents to exploit disease-specific molecular vulnerabilities (1,2). Nevertheless, diagnostic challenges remain and discordance rates of up to 60% between local and reference pathologists were recently reported in a study of >31,000 patients (3). Besides integrated clinical and phenotypic characterization, precise diagnosis often requires additional molecular and genetic analyses.

Cyclin D1 overexpression due to *CCND1* rearrangement (*CCND1*-R) involving immunoglobulin (IG) genes is considered a hallmark of mantle cell lymphoma (MCL). Recently, however, Cyclin D1 overexpression due to *CCND1*-R has also been identified in other B-cell lymphomas (4–6). Moreover, the morphological and phenotypic spectrum of MCL ranges from small-sized to large pleomorphic and blastoid cells, and negativity for CD5 and SOX11. In light of varying treatment strategies, distinguishing these cases accurately from other lymphomas poses an important diagnostic challenge.

Recently, unusual IGH class-switch recombination (CSR) or somatic hypermutation (SHM) was reported as an underlying mechanism for *CCND1*-R in such non-MCL lymphomas (7). To our knowledge, *in vitro* or *in vivo* model systems of these lymphomas are lacking (**Figure S1A, Supplementary Methods**). We report the establishment and comprehensive characterization of the *CCND1*-R non-MCL cell line *HaJo* and the corresponding patient-derived xenograft (PDX) model to close this gap and facilitate biological characterization and preclinical testing of novel therapies.

HaJo was derived from leukemic peripheral blood of a 74-year-old male patient initially diagnosed with splenic marginal zone lymphoma (SMZL) and managed with a watch- and-wait approach outside our center (**Figure 1A, Table S1, Supplementary Methods**). Upon disease progression, first-line treatment with six cycles of bendamustine and rituximab (BR) resulted in remission for six months. Subsequently, refractory disease with second-line ibrutinib treatment was observed and the patient was referred to our center. Here, diagnosis was revised to MCL based on *CCND1*-R (**Figure 1B**) with high Ki67 despite CD5- and SOX11-negativity (**Figure 1C**). The patient rapidly succumbed to his disease without receiving further treatment. Due to the unusual disease course and immunophenotype, we comprehensively characterized all available tumor samples by whole-exome sequencing (WES), whole-transcriptome sequencing (WTS), B-cell receptor sequencing (BCRseq) and targeted panel sequencing of IG and CSR regions (**Table S1, Supplementary Methods**).

**Figure 1:**
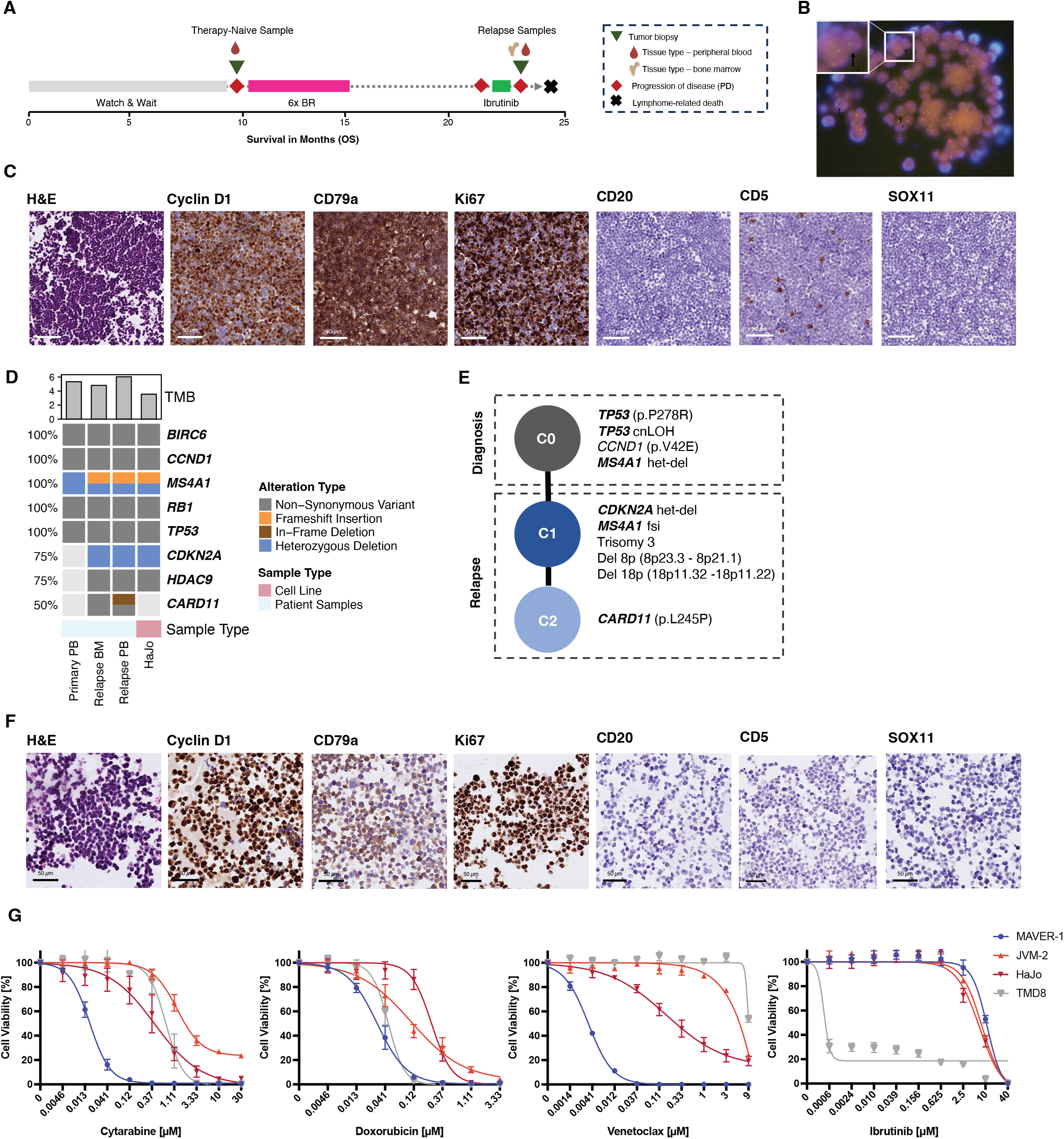
(A) The treatment-naive sample (green triangle) was collected from a 74-year-old male patient after disease progression (red diamond) under a watch-and-wait strategy. First-line therapy with six cycles of Bendamustine-Rituximab (BR) resulted in partial remission lasting six months, followed by disease relapse. The patient was subsequently treated with ibrutinib but exhibited refractory disease. At this time, two additional tumor samples were collected from peripheral blood and bone marrow. The patient ultimately succumbed to his disease. (B) FISH using a *CCND1* break-apart probe on a section of a cell block derived from unsorted leukemic PBMCs. Arrows indicate nuclei with split signals consistent with a *CCND1* rearrangement (78/100 nuclei counted). (C) Immunohistochemistry (IHC) staining of cell blocks generated from the patient’s peripheral blood at ∼40X magnification, which was utilized to establish the cell line which we termed HaJo acknowledging the contributions of <u>Ha</u>nnah Goldfarb-Wittkopf and <u>Jo</u>hanna Bihler. Malignant cells are of smaller size, with nuclei that exhibit slight irregularities and occasional indentations, while the chromatin appears somewhat dispersed. (D) Oncoprint summarizing key genetic driver alterations identified in the patient samples and the HaJo cell line. The bar chart above displays the tumor mutational burden (TMB), calculated as the number of non-synonymous variants per megabase (MB) of covered exome. (E) Tumor phylogeny depicting a linear clonal evolutionary pattern across primary patient samples collected before and after treatment. (F) Representative IHC stainings of cell block derived from the HaJo cell line at ∼40X magnification. (G) Relative cell viability of HaJo, the MCL cell lines MAVER-1 and JVM-2, as well as the ABC-DLBCL cell line TMD-8 serving as a positive control for ibrutinib sensitivity, assessed by CellTiter-Glo (CTG) after 72 hours of treatment with various concentrations of cytarabine, doxorubicin, venetoclax, and ibrutinib. Data represent the mean ± SD of at least three independent replicates.

Both therapy-naïve and relapsed lymphoma harbored a heterozygous deletion of *MS4A1* **(Figure 1D-E)**, as well as non-synonymous variants (NSV) in *BIRC6, CCND1, RB1* and *TP53*, including a copy-neutral loss of heterozygosity (cnLOH) affecting *TP53* **(Figure 1D,E)**. At relapse following BR, the tumor acquired NSVs in *HDAC9* and *CARD11*, a heterozygous deletion of *CDKN2A*, and an additional frameshift insertion in *MS4A1*. Phylogenetic reconstruction revealed a linear clonal trajectory, with *TP53* as the founding driver (C0), followed by a dominant clone (C1) harboring the *CDKN2A* deletion and biallelic *MS4A1* inactivation **(Figure 1E)**. The latter explains the complete loss of CD20 expression, potentially in response to selective pressure from rituximab **(Figure 1A,C,F)**. A subclone (C2) that emerged during ibrutinib treatment carried a known gain-of-function *CARD11* mutation (p.L245P) **(Figure 1D-E, Figure S1B)** that was lost in the HaJo cell line, potentially due to absence of therapeutic pressure (**Figure 1D**) (8). At the copy number level, we identified trisomy 3 and deletions involving short arms of chromosome 8 and 18 upon relapse **(Figure 1E, Figure S1C)**, both consistent with the complex karyotype **(Table S1)**. Trisomy 12 suggests a further layer of chromosomal complexity **(Table S1)**. Notably, structural abnormalities involving 9p13 cytoband, including t(9;10)(p13;q24) and add(9)(p13), suggesting disruption of both alleles at the *CDKN2A/B* locus.

To further delineate the mechanism of *IG*::*CCDN1* rearrangement, present in both the treatment-naïve and refractory sample (**Table S1**), we performed targeted panel sequencing covering all IG V(D)J and CSR regions as previously described (7). Despite this extensive panel, the rearrangement could not be detected by IgCaller, suggesting that the breakpoint resides within an intronic region, potentially altered by SHM. This rules out the canonical RAG-mediated, V(D)J anomalous rearrangement seen in most SOX11 positive and negative MCL (10,11) **(Table S4, Supplementary Methods)**. In line with these findings, germline identity of IG heavy chain variable region gene (IGHV) was 91,7% **(Table S4)**. Taken together, and in hindsight, the patient likely suffered from a transformation from a low-grade *CCND1*-R non-MCL. The patient’s rapid and ultimately fatal disease course (**Figure S1A**) underscores the critical unmet need for suitable model systems of *CCND1*-R non-MCL lymphomas.

To address this need, we first generated the HaJo cell line (**Figure S1A, Supplementary Methods**), mirroring the morphology (**Figure S1D**) and immunophenotype of the patient sample at relapse (**Figure 1C,F, Figure S1E-F**). Additionally, the molecular landscape of the patient sample is largely conserved in HaJo, both in terms of genetic alterations (**Figure 1D**) and transcriptional profiles (**Figure S1G**). Principal component (PC) analysis of transcriptome data revealed separation of SOX11-positive and SOX11-negative MCL cell lines along PC1, while HaJo clustered distinctly along PC2 (**Figure S1H**). To explore suitability of HaJo for *in vitro* studies, we treated HaJo, as well as MCL (SOX11-positive: MAVER-1, SOX11-negative: JVM-2) and non-MCL (TMD-8) cell lines with increasing doses of cytarabine, doxorubicin, venetoclax and ibrutinib (**Figure 1G**). While HaJo displayed an intermediate sensitivity to chemotherapy, we did not observe sensitivity of HaJo towards ibrutinib, thereby mimicking the resistance of the patient to ibrutinib **(Figure 1G)**. Intriguingly, we identified overexpression of BCL-2 in the patient (**Figure S1I**), and observed sensitivity to the BCL-2 inhibitor venetoclax **(Figure 1G)**.

Since such *in vitro* studies are inherently limited, we next explored the feasibility of generating a systemic PDX *in vivo* model. After intravenous injection of unsorted leukemic peripheral blood mononuclear cells (PBMCs) collected at disease progression on ibrutinib into NOD.Cg-*Prkdc*^*scid*^ *Il2rg*^*tm1Wjl*^/SzJ (NSG) mice **(Supplementary Methods)**, engraftment was observed. These F0 mice had to be sacrificed due to morbidity burden and enlarged spleens after a median of 138 days (range 78-167; **Figure 2A)**. Upon further intravenous passaging of tumor material isolated from spleens, median time to humane endpoint was <50 days in the F1 to F3 generation **(Figure 2A)**. Reflecting the systemic disease spread, all PDX mice showed infiltration of the spleen as assessed by immunohistochemistry (IHC) **(Figure 2B)**. While additional involvement of the bone marrow and/or liver was observed in most mice, leukemic disease was found in 25% (**Figure 2B**). Importantly, the PDX mimics the patient lymphoma and the cell line in terms of morphology and immunophenotype (**Figure 1C,F; Figure 2C**). Using IHC and flow cytometry, we were able to confirm a consistent immunophenotype across spleen, bone marrow, peripheral blood and liver of the PDX model (**Figure 2C,D**). In line with the molecular profiling of HaJo and the primary material, the genetic landscape and transcriptional signatures of the patient material were reflected by the PDX model across passages F0 to F3 (**Figure 2E & S2A**). To characterize the clonal architecture, we further performed BCRseq of the refractory patient sample, the HaJo cell line, and longitudinal passages of the PDX model. A dominant clone representing over 90% of the unique molecular identifier (UMI)-corrected reads was identified, embedded within a network of SHM **(Figure 2F)**, which was highly consistent across all of the different conditions **(Figure S2B)**.

**Figure 2:**
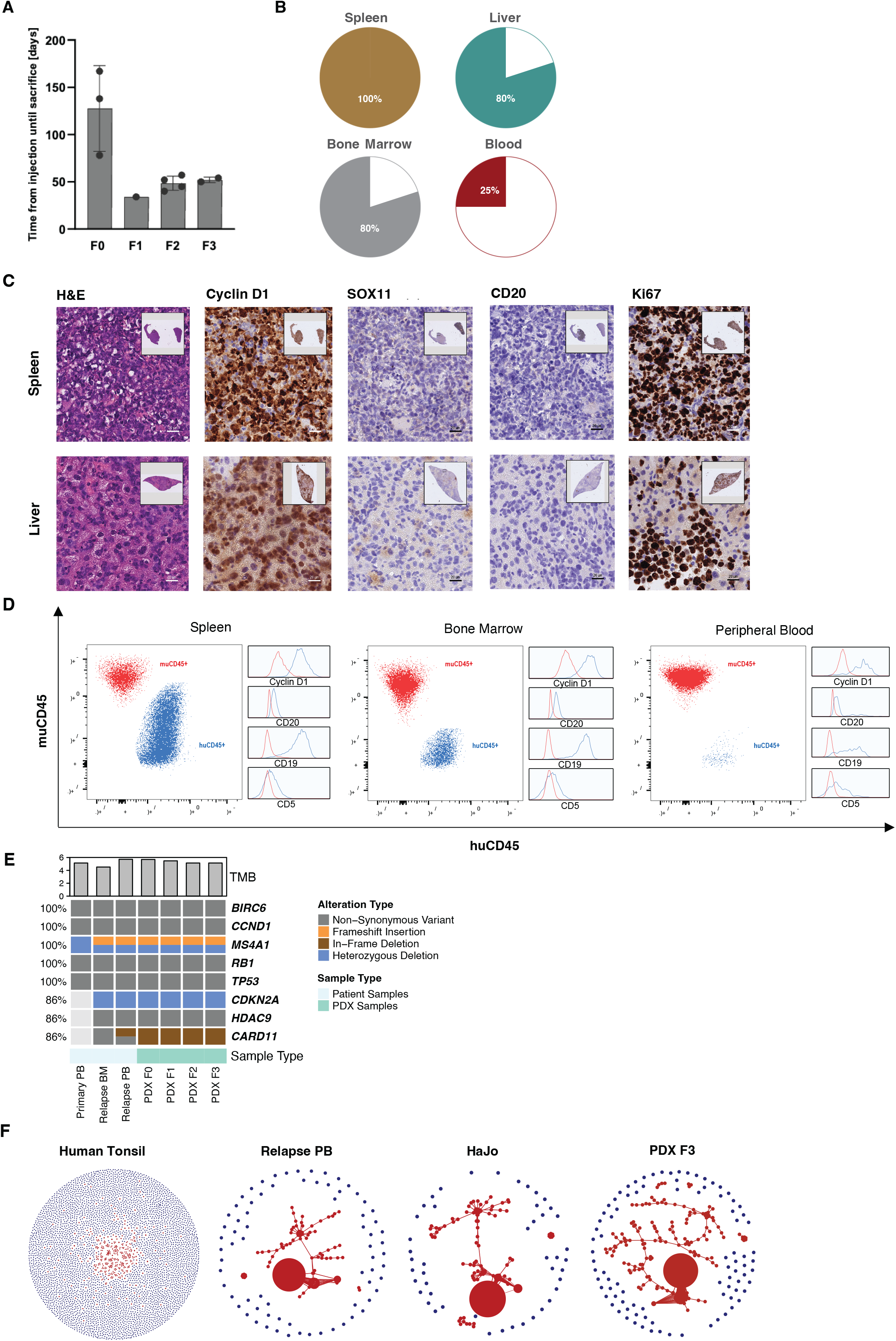
(A) Bar plot depicting time from intravenous tumor cell injection to death across PDX passages in days. (B) Relative fraction of organ infiltration as assessed by IHC staining. (C) Representative IHC images of spleen and liver from an F3 PDX mouse. H&E staining shows tumor cell infiltration in liver and spleen, with CCND1 and Ki67 overexpression, and SOX11 and CD20 negativity. (D) Flow cytometry plots showing murine CD45 and human CD45 expression, with additional depiction of expression of Cyclin D1, CD20, CD19 and CD5 in the corresponding populations by histograms in dissociated spleen tissue, bone marrow, and peripheral blood. (E) Oncoprint summarizing key genetic driver alterations identified in the primary patient sample and subsequent PDX passages. The bar chart above displays the tumor mutational burden (TMB), calculated as the number of non-synonymous variants per megabase (MB) of covered exome. (F) Clonality network plots derived from BCR sequencing data. Each circle represents a unique V(D)J sequence, and its size reflects the UMI-corrected read count. Clones which could be identified within an SHM tree are shown in red.

Our patient case and recent literature clearly illustrates that the presence of *CCND1*-R in these rare, aggressive non-MCL lymphomas present a significant diagnostic and therapeutic challenge (7). The conserved immunophenotype (CD19+, BCL2+, CD5-, CD10-, BCL6- and MUM1-) across patient material, HaJo cell line and corresponding PDX model shows striking similarities to the two previously reported transformed *CCND1*-R SMZL cases (7). Importantly, the genomic alterations identified at relapse, including trisomy 3 and 12 as well as deletions of *CDKN2A* and cytobands 8p and 18p, mirror changes recently described as characteristic of transformed SMZL (9). The successful generation of a systemic, multi-organ PDX model with high, fast and reproducible engraftment rates is a strength of our work. Unlike conventional cell line-derived xenografts or subcutaneous PDX models, commonly used in lymphoma research, our systemic model faithfully mimics systemic disease observed in human MCL patients and therefore offers a unique opportunity to explore fundamental questions surrounding the role of *CCND1*-R in non-MCL lymphomas. Functional experiments using our models could elucidate the contribution of *CCND1*-R to tumor proliferation and engraftment, thereby determining its therapeutic relevance in addition to its diagnostic value. A further notable feature of our model is the observed loss of CD20 expression caused by biallelic *MS4A1* inactivation, a phenomenon relevant for understanding and potentially overcoming resistance to anti-CD20 directed immunotherapies (12). Leveraging our systemic PDX in a humanized *in vivo* background allows to evaluate the efficacy of immunotherapeutic approaches in the context of CD20 loss. Moreover, consistent overexpression of BCL2 in our model supports the rationale to explore BCL2 inhibition as a targeted therapeutic strategy.

Nonetheless, additional models representing the molecular heterogeneity of *CCND1*-R non-MCL are needed to comprehensively characterize this entity. These will be essential to differentiate between subtype-specific vulnerabilities and to refine diagnostic and therapeutic strategies.

Taken together, both the patient-derived HaJo cell line and corresponding systemic PDX *in vivo* model mimic key features of SOX11-negative *CCND1*-R non-MCL and thereby constitute valuable tools to study these rare but challenging lymphomas.

## Software

All statistical analyses and visualizations were conducted using R (version 4.3.2) or GraphPad Prism (version 10.4.2). Heatmaps and oncoprints were generated with ComplexHeatmap package (version 2.22.0) or pheatmap package (version 1.0.12).

## Acknowledgments

We are grateful to the patient and his family. We further thank Hannah Goldfarb-Wittkopf, Mathilde De Jong and Matthew Arnett for excellent technical assistance. This work was supported by the Max Planck Biology for Ageing Core Facilities: All flow cytometry analyses were performed in the FACS&Imaging Core Facility. Breeding of mice and husbandry were performed in the Comparative Biology Core Facility and supported by Sabrina Seyfarth, Frederik Böddeker, Jerome Henn and Bettina Bertalan. We thank Alexandra Florin and Marion Müller from the Pathology department of the University Hospital Cologne for support with tissue embedding and immunohistochemistry stainings. This work was supported by the DFG Research Infrastructure West German Genome Center (project 407493903) as part of the Next Generation Sequencing Competence Network (project 423957469). NGS analyses were carried out at the production site Cologne (Cologne Center for Genomics (CCG)). This work was funded by the Max Planck Society (to R.D.J), the German Research Foundation (grant no. 496650118, 5504 FOR JA 2439/5-1 TP05 to R.D.J., JA2439/4-1 to R.D.J., grant 455784452 as part of CRC1530, with project funding for R.D.J. (C02), the Ministry for Culture and Science North-Rhine-Westphalia (NW21-062A CANTAR to R.D.J.), the Behrens-Weise-Foundation (Grant for the Improvement of Human Health to R.D.J), the Center for Molecular Medicine Cologne (Project A10 to R.D.J.), the German Cancer Aid (Post-doctoral Fellowship Stipend by the *Mildred Scheel School of Oncology* to P.J.B.), the Else Kröner-Fresenius-Foundation (Excellence Stipend to P.J.B.), the Else Kröner Forschungskolleg Clonal Evolution in Cancer (C.M.G) and the KölnFortune Clinician Scientist Program of the University of Cologne (M.R.).

## Author Contributions

P.J.B. and R.D.J. conceived the study. J.B., P.J.B. collected patient samples and established the cell line. C.M.G., A.V.L., N.M., C.B., M.L., M.T., P.J.B. established PDX models and performed mouse experiments. A.V.L., N.M., C.B., P.J.B. performed FACS experiments and analyzed data. C.M.G., S.B. analyzed and interpreted genomic and transcriptomic data. E.C., F.N. performed the analysis and interpretation of the targeted sequencing data. J.M. reviewed pathology. M.R. performed drug assays. M.T., P.J.B., R.D.J. supervised the study. C.M.G., P.J.B., R.D.J. wrote the first draft of the manuscript. All authors reviewed, revised and approved the final version of the manuscript.

## Data Availability Statement

The HaJo cell line and early passages of the HaJo PDX model as well as the data generated in this study are available from the corresponding author upon reasonable request.

## Figure Legends

<u>Supplementary Figure 1</u>

(A) Distribution of *CCND1* expression across cell lines of B-cell malignancy. Log_2_(TPM+1) values are shown for 142 DepMap B-Cell Non-Hodgkin Lymphoma cell lines grouped by primary disease category. Density ridges depict the distribution of expression levels within each group. Individual cell lines are overlaid as dots; selected cell lines with high *CCND1* expression (>6.5) and known annotation as either Mantle Cell Lymphoma (blue), Multiple Myeloma (yellow) are highlighted and labeled. The red dot denotes HaJo, a SOX11-negative *CCND1*-R cell line sequenced independently. Vertical dashed lines indicate group-wise medians.

(B) Clonal composition of each sample, illustrating changes in clonal dynamics over time.

(C) Copy number profiles of the primary PB sample (purity: 0.86, ploidy: 2.00), relapse PB sample (purity: 0.68, ploidy: 2.08) and HaJo (purity: 0.93, ploidy 2.05). Total copy number is shown in brown, and minor allele copy number in green.

(D) Brightfield microscopy images of HaJo cells at 4X and 20X magnification.

(E) Overview of IHC staining of cell block generated from the patient’s peripheral blood at time of relapse, which was utilized to establish the HaJo cell line.

(F) Overview of IHC stainings of cell block derived from the HaJo cell line.

(G) GSVA scores from ssGSEA comparing patient material and the HaJo cell line.

(H) PCA of the 500 most variable genes comparing the HaJo cell line to SOX11-positive (blue) and SOX11-negative (yellow) cell lines.

(I) Representative IHC stainings from the patient’s bone marrow at relapse, shown at low (∼4X, top row) and high (∼40X, bottom row) magnification.

<u>Supplementary Figure 2</u>

(A) GSVA scores from ssGSEA comparing patient material at time of relapse with PDX passages.

(B) Read fraction of B-cell receptor (BCR) clones derived from the heavy-chain library, showing consistent BCR clonotype across the primary patient material, PDX generations, and the HaJo cell line.

## References

1. Zhang MC, Tian S, Fu D, Wang L, Cheng S, Yi HM, et al. Genetic subtype-guided immunochemotherapy in diffuse large B cell lymphoma: The randomized GUIDANCE-01 trial. Cancer Cell. 2023 Oct;41(10):1705-1716.e5.

2. Palmer AC, Kurtz DM, Alizadeh AA. Cell-of-Origin Subtypes and Therapeutic Benefit from Polatuzumab Vedotin. N Engl J Med. 2023 Aug 24;389(8):764–6.

3. Syrykh C, Chaouat C, Poullot E, Amara N, Fataccioli V, Parrens M, et al. Lymph node excisions provide more precise lymphoma diagnoses than core biopsies: a French Lymphopath network survey. Blood. 2022 Dec 15;140(24):2573–83.

4. Yoshida M, Ichikawa A, Miyoshi H, Kiyasu J, Kimura Y, Arakawa F, et al. Clinicopathological features of double‐hit B ‐cell lymphomas with MYC and BCL 2, BCL 6 or CCND 1 rearrangements. Pathol Int. 2015 Oct;65(10):519–27.

5. Nishida Y, Takeuchi K, Tsuda K, Ugai T, Sugihara H, Yamakura M, et al. Acquisition of t(11;14) in a patient with chronic lymphocytic leukemia carrying both t(14;19)(q32;q13.1) and +12. Eur J Haematol. 2013 Aug;91(2):179–82.

6. Koduru PR, Chen W, Garcia R, Fuda F. Acquisition of a t(11;14)(q13;q32) in clonal evolution in a follicular lymphoma with a t(14;18)(q32;q21) and t(3;22)(q27;q11.2). Cancer Genet. 2015 Jun;208(6):303–9.

7. Özoğul E, Montaner A, Pol M, Frigola G, Balagué O, Syrykh C, et al. Large B-cell lymphomas with CCND1 rearrangement have different immunoglobulin gene breakpoints and genomic profile than mantle cell lymphoma. Blood Cancer J. 2024 Sep 23;14(1):166.

8. Stinson JR, Dorjbal B, McDaniel DP, David L, Wu H, Snow AL. Gain-of-function mutations in CARD11 promote enhanced aggregation and idiosyncratic signalosome assembly. Cell Immunol. 2020 Jul;353:104129.

9. Grau M, López C, Navarro A, Frigola G, Nadeu F, Clot G, et al. Unraveling the genetics of transformed splenic marginal zone lymphoma. Blood Adv. 2023 Jul 25;7(14):3695– 709.

10. Nadeu F, Mas-de-les-Valls R, Navarro A, Royo R, Martín S, Villamor N, et al. IgCaller for reconstructing immunoglobulin gene rearrangements and oncogenic translocations from whole-genome sequencing in lymphoid neoplasms. Nat Commun. 2020 Jul 7;11(1):3390.

11. Nadeu F, Martin-Garcia D, Clot G, Díaz-Navarro A, Duran-Ferrer M, Navarro A, et al. Genomic and epigenomic insights into the origin, pathogenesis, and clinical behavior of mantle cell lymphoma subtypes. Blood. 2020 Sep 17;136(12):1419–32.

12. Rushton CK, Arthur SE, Alcaide M, Cheung M, Jiang A, Coyle KM, et al. Genetic and evolutionary patterns of treatment resistance in relapsed B-cell lymphoma. Blood Adv. 2020 Jul 14;4(13):2886–98.

13. Cun Y, Yang TP, Achter V, Lang U, Peifer M. Copy-number analysis and inference of subclonal populations in cancer genomes using Sclust. Nat Protoc. 2018 Jun;13(6):1488– 501.

14. Peifer M, Fernández-Cuesta L, Sos ML, George J, Seidel D, Kasper LH, et al. Integrative genome analyses identify key somatic driver mutations of small-cell lung cancer. Nat Genet. 2012 Oct;44(10):1104–10.

15. George J, Lim JS, Jang SJ, Cun Y, Ozretić L, Kong G, et al. Comprehensive genomic profiles of small cell lung cancer. Nature. 2015 Aug 6;524(7563):47–53.

