## Supplementary Methods for "Establishment and Characterization of a *CCND1*-Rearranged Non-Mantle Cell Lymphoma Cell Line and Patient-Derived Xenograft Model"

**Supplementary Material**

**Materials & Methods**

**Generation of HaJo Cell Line and Patient-Derived Xenograft (PDX) Model**

Written informed consent was obtained from the patient and analysis of patient material conducted in accordance with approvals by the institutional review board of the University of Cologne (20-1577, 21-1472). For cell line and PDX generation, mononuclear cells were isolated by Histopaque density gradient (Sigma Aldrich, 10771) from fresh peripheral blood of the patient according to manufacturer’s instructions. For generation of the HaJo cell line, fresh unsorted PBMCs with a reported MCL cell content of 39% were plated at an initial concentration of 1x10^6^/mL in RPMI-1640 medium (Thermo Fisher, 21875091) + 10% FCS (Thermo Fisher, A5256701) + P/S (Thermo Fisher, 15070063) and monitored regularly. After approximately 8 weeks, expansion of the malignant B-cell population was observed and cells passaged 2-3x/week to maintain concentrations of 0.5-1.5 x 10^6^/mL at a doubling time of approximately 40 hours.

For all *in vivo* experiments, 4- to 6-week-old NSG mice obtained from Charles River Laboratories were used. The general health status of all animals was monitored daily. All experiments were performed in accordance with the guidelines of the Federation of European Laboratory Animal Science Associations (FELASA), approved by the respective regional authorities (LANUV: 81-02.04.2023.A091) and were conducted in the animal facilities of the Max-Planck Institute for Biology of Ageing, Cologne, Germany.

For PDX generation, unsorted PBMCs were thawed and washed twice with PBS. A total of 20 x 10^6^ cells were kept on ice until immediate transplantation via tail vein injection into NSG mice (F0 generation). Mice were sacrificed upon reaching a humane endpoint due to morbidity burden. At the time of sacrifice, spleen, bone marrow, liver, peripheral blood and all macroscopically affected organs were collected. Depending on sample type, amount of tissue and desired downstream applications, samples were stored either formalin-fixed paraffin-embedded (FFPE), snap frozen in liquid nitrogen and/or as viable single-cell suspensions (scSuspension) after mechanic dissociation with a 100µM cell-strainer (Falcon, 352360). Parts of the spleen tissue were passaged further by tail vein injection after dissociation: for the F1 generation, 10 x 10^6^ cells per mouse were injected while 1 x 10^6^ cells per mouse were injected for the subsequent F2 and F3 generations from intermittently frozen spleen scSuspensions.

**FISH**

To visualize the translocation of *CCND1* in the refractory patient sample, FISH was performed on interphase nuclei using a dual-color break apart probe for 11q13.3 (Cat. Z-2108-200, Zytovision, Germany) according to the manufacturer’s protocol. Signal patterns were evaluated manually on a Leica DM 5500B, and if not stated otherwise 100 translocation-positive nuclei were quantified. All other FISH results, including t(11;14), were obtained by the diagnostic laboratories as part of analyses in routine clinical care with validated assays.

**RNAseq**

Bulk RNAseq data of 17 samples were generated, comprising both cell lines and primary/patient-derived xenograft (PDX) tissues. Cell lines were obtained from publicly available repositories (DSMZ, ATCC) or collaborators (Prof. Dreyling, Munich; Prof. Klapper, Kiel; Dr. Zhao, Cleveland) and were cultured under optimized conditions using either RPMI-1640 supplemented with 10–20% FCS or DMEM (Gibco, 61965059; for Granta cell line). All cell lines were authenticated by short tandem repeat (STR) profiling and routinely tested to exclude mycoplasma contamination. RNA was extracted using the RNeasy Mini Kit (Qiagen, 74106) according to the manufacturer’s protocol. The RNA integrity was assessed with the Agilent TapeStation using the RNA ScreenTape. RNA libraries from cell lines were prepared using 500ng of total RNA with the Illumina TruSeq Strand kit including ERCC spike-ins, while libraries from primary and PDX samples were prepared from 10ng total RNA using the SMART-Seq Stranded Kit (Takara Bio, 20020595) enabling total RNA sequencing. Libraries were sequenced either on the Illumina NovaSeq X Plus platform, generating 2 × 101 bp paired-end reads or on an Illumina NovaSeq6000 sequencing instrument with a PE100 protocol aiming for 50 million clusters. The resulting data were processed using the nf-core/rnaseq pipeline (version 3.14.0) executed with Nextflow (version 23.10.1), aligning reads to the human reference genome (GRCh38, Ensembl release 109) using STAR (version 2.7.9a). Quality assessment of raw reads was performed with FastQC (version 0.12.1), and adapter trimming was conducted using Trim Galore (version 0.6.7) incorporating Cutadapt (version 3.4). Transcript quantification was achieved with Salmon (version 1.10.1), and gene-level counts were generated using tximport (Bioconductor-tximeta version 1.12.0). Quality control metrics were compiled using RSeQC (version 5.0.2), Qualimap (version 2.3), and dupRadar (version 1.28.0). Duplicate reads were marked with Picard (version 3.0.0), and comprehensive quality metrics were summarized using MultiQC (version 1.19). Differential gene expression analysis was conducted using DESeq2 (version 1.48.0) within the R environment (version 4.3.2). To assess pathway activity at the individual sample level, single-sample gene set enrichment analysis (ssGSEA) was performed using the GSVA package (version 3.21) in R. Normalized expression data were analyzed with a predefined gene set collection using ssgseaParam() and gsva().

**Whole-Exome Sequencing**

We performed whole-exome sequencing for the cell line with the SureSelect Human All Exon V6 Kit (Agilent) and for patient and PDX samples with the Twist Human Core Exome kit with RefSeq and Mitochondrial Panel (Twist Bioscience), following the protocol of the manufacturer. Exon-enriched libraries were subjected to paired-end sequencing on either the Illumina NovaSeq platform. Libraries were prepared to reach a mean insert size of 200 base pairs (bp) for sequencing with a read length of 2× 100 bp. Tumor DNA material was sequenced aiming for a coverage of at least 200× which, following filtering of PCR-duplicated reads and alignment to the annotated human genome (hg19), resulted in an average coverage of 214X. Patient tumor samples showed a median purity of 82%, thus minimizing problems in the assessment of tumor-specific mutations. This allowed for sufficient sequencing depth for reliable analysis for allelic fractions and clonality, as described below.

**Data processing of Whole-Exome Sequencing**

Raw sequencing reads were processed as previously described (13–15). Reads were aligned to the human reference genome (GRCh37/hg19). For PDX tumor samples, sequencing reads were aligned to a combined human and murine reference genome (GRCh37/hg19 and GRCm38/mm10), to exclude sequencing reads from murine cells and to allow for uniform processing of all samples from a given patient. Concordant read-pairs were identified as potential PCR duplicates and were subsequently masked in the alignment file and annotated as the number of masked reads.

Human sequencing reads (mapped to the human reference genome) were analyzed for tumor purity, tumor ploidy, somatic mutations and copy number alterations. Mutation calling was performed as previously described (14,15). In brief, variant counts were assessed for tumor and matching normal samples, corrected for sequencing noise and compared with a database of 300 whole-exome and genome sequenced normal samples to filter and determine somatic mutation calls. Variants at low allelic fractions are often prone to result from sequencing artefacts, which occur as a consequence of sequencing noise arising from high-coverage WES due to either fragmented DNA as part of FFPE material or low-level contamination with murine reads in tumors derived from murine xenograft models. We therefore implemented strict filtering criteria for mutations occurring at allelic fractions of less than 0.2. Mutations were then filtered out if (1) the forward–reverse score was below 0.2 (forward–reverse score is 1.0 if 50% of variant reads are found on the forward or reverse read, and 0 if all variant reads are on one orientation); and (2) the allelic fraction of the variant *v* in consideration of minimal coverage *C* of the normal or matching tumor sample at position *i* (*C_i_*^min(tumour/normal)^) did not exceed the read count (rc) threshold with a default value of 15. This was calculated as *C_i_*^min(tumour/normal)^ × *v_i_* < rc. We thus introduced a decision boundary that filters out mutations at relatively low allelic fractions and low sequencing coverage; mutations with low allelic fractions but high coverage were retained for further analyses. In addition, we adjusted the stringency of this cut-off for individual samples. Although this stringent cut-off limits the identification of subclonal mutations, we have thus controlled for potential sequencing noise and false-positive mutation calls. As described below, longitudinal studies may suggest mutations at very low allele fractions in one tumor that might be more abundant at another tumor site. In this instance, truly subclonal mutations at low allelic fractions that were filtered out in one sample at this step of the analysis were reintroduced as somatic mutation calls if the same mutation passed all stringent filtering criteria in another matched tumor sample.

**Analysis of tumor phylogenies**

We inferred cancer cell fractions (CCFs) by comparing observed to expected allele frequencies, accounting for tumor purity, ploidy, and local copy number. Mutations were clustered based on their CCFs to infer subclones, and two-dimensional clustering was applied to paired samples to reconstruct clonal evolution as previously described (15). Shared mutations, subclonal architecture, and phylogenetic relationships were determined using this approach, with additional filters for mapping quality, loss-of-heterozygosity, and low-powered mutation calls.

### **Custom IG panel sequencing and bioinformatic analysis**

To characterize the IG gene rearrangements and identify the IG breakpoint regions and *CCND1*-R, we used a previously described, customized capture-based NGS panel (IG-MCL panel) covering the full-length IGH V(D)J regions (from FR1 to FR4), IGK and IGL VJ, and CSR (7). Bioinformatics analysis was performed as previously described (7). Briefly, raw reads were trimmed using Trimmomatic (version 0.40) and aligned to the GRCh38 reference human genome using the BWA-mem algorithm (version 0.7.17). PCR and optical duplicates were marked using MarkDuplicates from Picard (version 2.24.0,<https://broadinstitute.github.io/picard/>). IG gene rearrangements, identity, and *IG::CCND1* rearrangement breakpoints were analyzed using IgCaller (version 1.4) (10).

### **B-Cell Receptor Sequencing (BCRseq)**

Total RNA was isolated from primary tumor material, patient-derived xenografts (PDXs), and the HaJo cell line as described above. RNA quality and integrity were assessed using the Agilent TapeStation system. Libraries were prepared using the SMARTer Human BCR IgG/IgM/IgA profiling kit (Takara Bio, 634776) with integrated unique molecular identifiers (UMIs), following the manufacturer’s protocol. Paired-end sequencing (2 × 300 bp) was performed on an Illumina MiSeq platform (MiSeq Reagent Kit v3, 600 cycles) to ensure full-length coverage of the immunoglobulin heavy chain transcripts.

Raw sequencing data were processed using MiXCR (version 4.7.0) for alignment, clonotype assembly, and UMI-based error correction. Clonotypes were assembled using the full-length VDJRegion. Unique molecular identifiers (UMIs) were used for error correction and quantification. Clonotypes were defined based on identical VDJRegion nucleotide sequences and consistent V and J gene usage. SHM lineage trees were reconstructed using MiXCR’s findAlleles and findShmTrees module. From these SHM trees, network edges were derived by identifying pairwise shortest paths connecting clonotypes within a tree.

For visualization, networks were generated using Gephi (version 0.10.1). In Gephi, node size was scaled according to UMI-based clone abundance (uniqueMoleculeFraction), and edges reflected inferred mutational relationships within SHM trees. Downsampling was applied where necessary to reduce dataset complexity.

**Immunohistochemical (IHC) analysis**

Fresh tissue was fixed in 4% paraformaldehyde (PFA) for 24 hours and subsequently embedded in paraffin. Sections of 4μm were stained with antibodies listed in **Table S2**.

**Flow cytometry**

Flow cytometry was performed using a LSRFortessa cell analyzer (BD Biosciences) and data was analyzed using the FlowJo software (version 10.6.1; BD Biosciences). Single-cell suspensions from mouse spleen, bone marrow or peripheral blood were collected after mechanical dissociation (100µM cell-strainer; Falcon, 352360) and preserved at -80°C in FCS+10% DMSO until staining with fluorescent dye-labeled antibodies (**Table S3**). Permeabilization for intracellular staining was performed using the eBioscience FoxP3/Transcription Factor Staining Buffer Set (Invitrogen, 00-5523-00).

**Drug Assays**

Cell viability was assessed by CellTiter-Glo® (Promega, G7572) and in line with the manufacturer’s instructions. All experiments were conducted for a minimum of three replicates. In summary, 10.000 cells (MAVER-1, TMD8) or 15.000 cells (JVM-2, HaJo) per well were seeded on a 96-well plate (ThermoFisher, 236105) in triplicates. Cytarabine (Selleckchem, S1648), Doxorubicin (MedChemExpress, HY-15142), Venetoclax (MedChemExpress, HY-15531), and Ibrutinib (MedChemExpress, HY-10997) were diluted to the desired concentrations in RPMI and added to the designated wells for a total volume of 100 µL with DMSO serving as control. After incubation for 72h, CellTiterGlo reagent was added to each well. Following incubation on an orbital shaker for 15 minutes, luminescence was measured on a Tecan Spark Microplate Reader (Tecan).

**Supplementary Tables**

Supplementary Table 1: Summarized pathology reports

| **Tissue** | **Type** | **Timepoint** | **Positive** | **Negative** |
| --- | --- | --- | --- | --- |
| Peripheral Blood | FACS | Therapy-  naive (after watch&wait) | 75% B-lymphocytes, CD19, CD20, CD22, CD79b, FMC7 | CD5, CD10, CD23 |
| Peripheral Blood | Cyto-genetics | Therapy-  naive (after watch&wait) | t(11;14)(q13;q32) |  |
| Bone Marrow | IHC | Relapse | CD79a, Cyclin D1, PAX5, DBA44part,  p53, Ki67 (50%), MYC, BCL2 | CD5, CD20, CD23, MUM1, SOX11, CD123, BCL6  BRAF, TdT, LEF1, CD38, CD10, Annexin A1, CD11c |
| Bone Marrow | Molecular Genetics | Relapse | *TP53*, Exon 8, c.833C>G, p.(P278R), AF 72%, Coverage 4483x | *BAX, BCL2, BRAF, BTK, CXCR4, EZH2, KIT,*  *KRAS, MCL1-, MYD88, NOTCH1, NRAS, PLCG2* |
| Bone Marrow | Cyto-genetics | Relapse | Karyotype: 47,X,-Y,+3,t(6;12)(q25;q13),der(8;18)(q10;p10),t(9;10)(p?13;q?24),add(9)(p13),-11,t(11;14)(q13;q32),+12,der(12)add(12)(p11)add(12)(q11),der(17)t(11;17)(q13;q25),+2mar[11]/46,XY[1].nuc ish (BCL6x3)[73/132], ],(MYCx2,IGHx3)[55/109],(PCM1,JAK2)x1[54/119,(ATM,TP53)x2[107],(12cenx3,DLEUx2,LAMP1x2)[42/104],(IGHx3,BCL2x2)[47/113] |  |
| Bone Marrow | FACS | Relapse | 60% lymphocytes, 54% B cells of leukocytes, CD19 (dim), CD79b, FMC7, IgM, Kappa | CD5, CD10, CD3 |
| Peripheral Blood | FACS | Relapse | 46% Lymphocytes, 39% B cells of leukocytes, CD19 (dim), CD79b, FMC7, IgM, Kappa | CD5, CD10, CD3 |

Supplementary Table 2: List of IHC antibodies used.

| **Antigen** | **Clone** | **Dilution** | **Pre-handling** | **Manufacturer** |
| --- | --- | --- | --- | --- |
| CD20 | L26 | 1:1250 | Citrat | Dako |
| CD5 | 4C7 | 1:100 | EDTA | Cellmarque |
| Cyclin D1 | QR002 | 1:200 | EDTA | Quartett |
| Ki67 | MIB1 | 1:200 | EDTA | Zeta |
| SOX11 | ZSX11 | 1:50 | Citrat | Zytomed |
| CD79a | JCB117 | 1:800 | Citrat | Dako |

Supplementary Table 3: List of flow cytometry antibodies used.

| **Antigen** | **Conjugate** | **Clone** | **Dilution** | **Manufacturer** | **Identifier** |
| --- | --- | --- | --- | --- | --- |
| Fixable Green Live/Dead | FITC |  | 1:2000 | Thermo Fisher | L23101 |
| Zombie NIR | APC/Fire750 |  | 1:2000 | BioLegend | 423105 |
| CD45 (anti-mouse) | PECy7 | 30-F11 | 1:500 | BioLegend | 103113 |
| CD45 (anti-human) | BV785 | HI30 | 1:100 | BioLegend | 304047 |
| CD5 (anti-human) | PerCP-Cy5.5 | L17F12 | 1:100 | BioLegend | 364005 |
| Cyclin D1 (anti-human) | PECy5 | SP4 | 1:1000 | Abcam | ab16663 |
| CD20 (anti-human) | BV510 | 2H7 | 1:100 | BioLegend | 302340 |
| CD19 (anti-human) | BUV737 | SJ25C1 | 1:200 | BD Biosciences | 612757 |
| Human Fc Block |  | Fc1 | 1:200 | BD Biosciences | 564220 |
| Mouse Fc Block |  | 2.4G2 | 1:200 | BD Biosciences | 553141 |
