## Supplementary figures and images for "Establishment and Characterization of a *CCND1*-Rearranged Non-Mantle Cell Lymphoma Cell Line and Patient-Derived Xenograft Model"

### Supplementary Figures

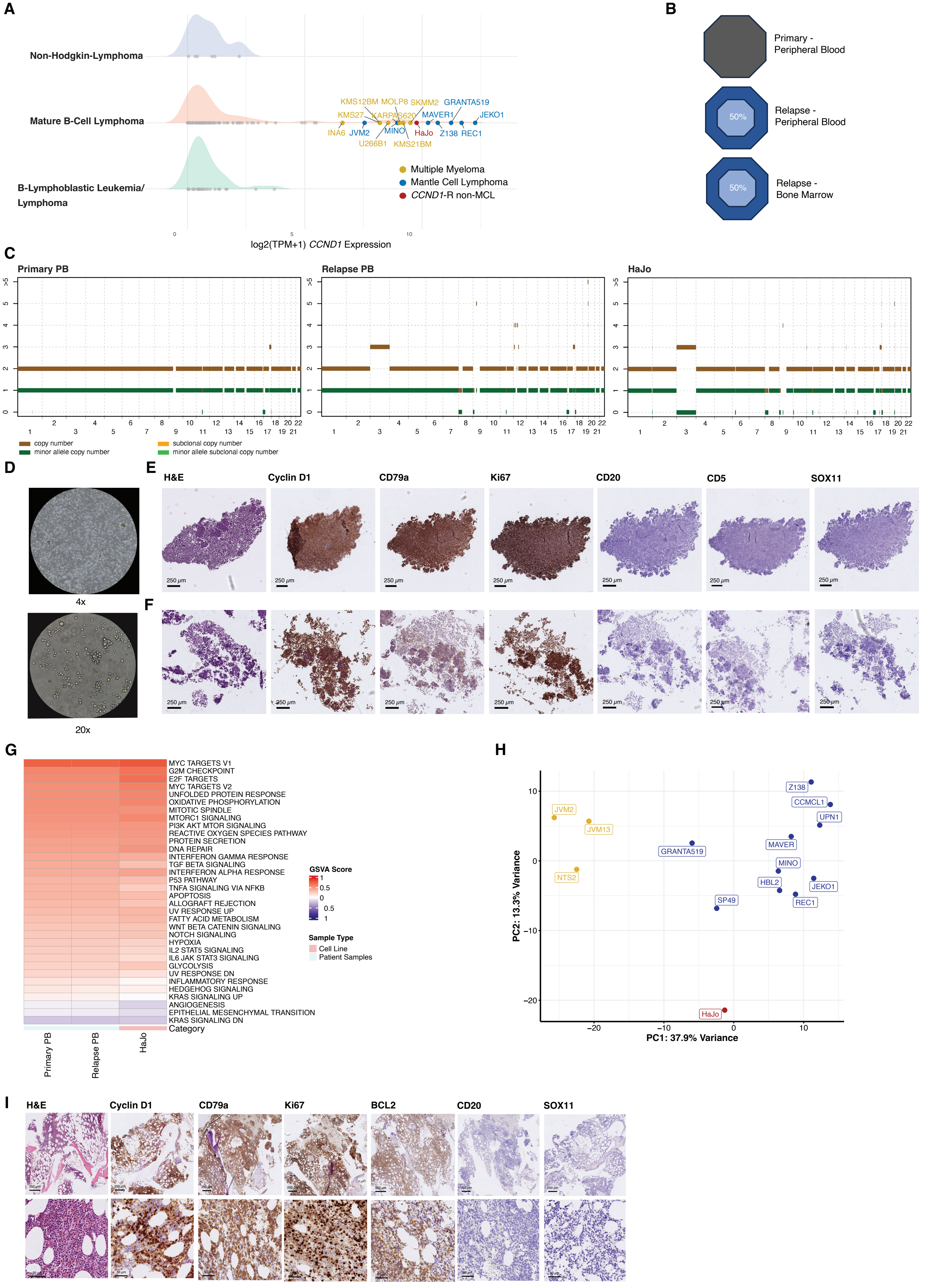


Supplementary Figure 1

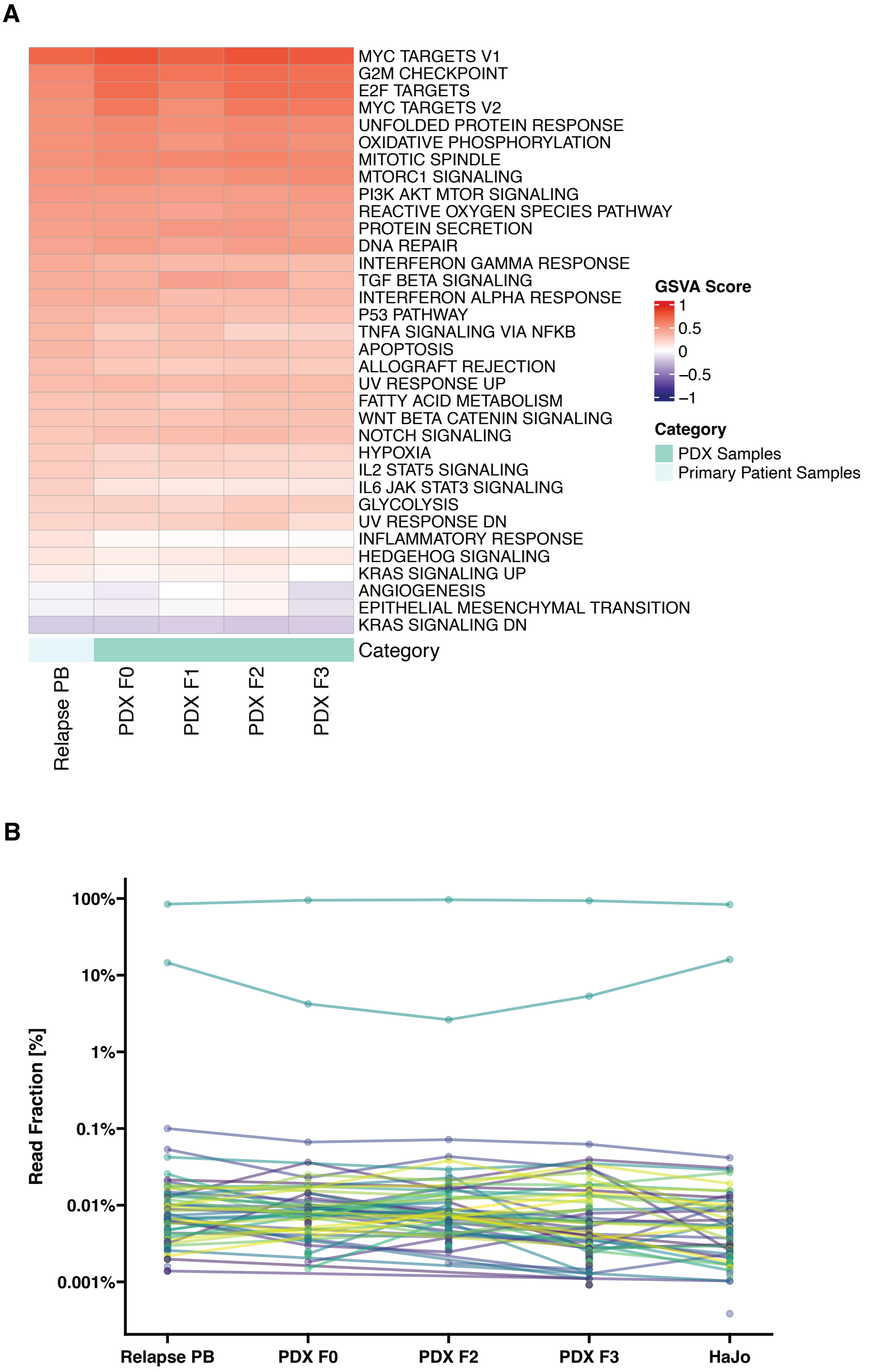


Supplementary Figure 2
